# Endocannabinoid regulation of inward rectifier potassium (Kir) channels

**DOI:** 10.1101/2024.05.29.596143

**Authors:** Sultan Mayar, Mariia Borbuliak, Andreas Zoumpoulakis, Tahar Bouceba, Madeleine M. Labonté, Ameneh Ahrari, Niveny Sinniah, Mina Memarpoor-Yazdi, Catherine Vénien-Bryan, D. Peter Tieleman, Nazzareno D’Avanzo

## Abstract

The inward rectifier potassium channel Kir2.1 (KCNJ2) is an important regulator of resting membrane potential in both excitable and non-excitable cells. The functions of Kir2.1 channels are dependent on their lipid environment, including the availability of PI(4,5)P_2_, secondary anionic lipids, cholesterol and long-chain fatty acids acyl coenzyme A (LC-CoA). Endocannabinoids are a class of lipids that are naturally expressed in a variety of cells, including cardiac, neuronal, and immune cells. While these lipids are identified as ligands for cannabinoid receptors (CBRs), there is a growing body of evidence that they can directly regulate the function of numerous ion channels independently of CBRs. Here we examine the effects of a panel of endocannabinoids on Kir2.1 function and demonstrate that a subset of endocannabinoids can alter Kir2.1 conductance to varying degrees independently of CBRs. Using computational and SPR analysis, endocannabinoid regulation of Kir2.1 channels appears to be the result of altered membrane properties, rather than through direct protein-lipid interactions. Furthermore, differences in endocannabinoid effects on Kir4.1 and Kir7.1 channels, indicating that endocannabinoid regulation is not conserved among Kir family members. These findings may have broader implications on the function of cardiac, neuronal and/or immune cells.

## INTRODUCTION

Cannabinoids are a special class of lipids, largely identified by their ability to activate CB1 and CB2 cannabinoid receptors (CBRs) (Howlett, 2002). They are typically defined by their source, with phytocannabinoids present in plants, endocannabinoids endogenously expressed in various mammalian cells, and synthetic cannabinoids manufactured exogenously. Endocannabinoids are part of the endocannabinoid system that regulates the signalling of a variety of biological activities. Additionally, endocannabinoid-like molecules, which are chemically related and often byproducts but may not activate CBRs, also have important biological functions, including energy homeostasis and metabolic regulation. In addition to regulating ion channel function via changes in intracellular signalling (Maroso et al., 2016), cannabinoids also regulate various ion channels independently of CBRs. This includes members of the TRP family (De Petrocellis & Di Marzo, 2011; De Petrocellis et al., 2012; Iannotti, Silvestri, et al., 2014), Nav channels (Ghovanloo et al., 2021; Ghovanloo et al., 2018; Okada et al., 2005), Cav channels (Ross, Napier, & Connor, 2008), Kv channel (Iannotti, Hill, et al., 2014), HCN channels (Mayar, Memarpoor-Yazdi, Makky, Eslami Sarokhalil, & D’Avanzo, 2022; Page & Ruben, 2024), and Gly receptors (Ahrens et al., 2009; Hejazi et al., 2006) among others.

Inward rectifier potassium (Kir) channels selectively control the permeation of K^+^ ions across cell membranes, controlling a variety of cellular functions including maintaining resting membrane potential, modulation of cellular excitability, and regulation of whole-body electrolyte homeostasis. Members of this ion channel family are highly sensitive to regulation the lipids in which they are embedded. For example, Kir channels are directly regulated by phosphoinositides (PIPs) in the absence of other proteins or downstream signaling pathways (W. W. Cheng, Enkvetchakul, & Nichols, 2009; D’Avanzo, Cheng, Doyle, & Nichols, 2010; Enkvetchakul, Jeliazkova, & Nichols, 2005; Leal-Pinto et al., 2010). Additionally, Kirs are regulated by other lipids including anionic phospholipids, cholesterol, long chain co-A (W. W. Cheng et al., 2009; W. W. L. Cheng, D’Avanzo, Doyle, & Nichols, 2011; D’Avanzo et al., 2010; D’Avanzo, Hyrc, Enkvetchakul, Covey, & Nichols, 2011; Enkvetchakul et al., 2005; Furst, Nichols, Lamoureux, & D’Avanzo, 2014; Rosenhouse-Dantsker, Noskov, Logothetis, & Levitan, 2013; Singh, Shentu, Enkvetchakul, & Levitan, 2011). Some evidence suggests that K_ATP_ channels (Kir6.x channels with their partnering sulfonyl urea receptor (SUR)) are modified by endocannabinoids anandamide (AEA) and 2-Arachidonoyl Glycerol (2-AG) (Li et al., 2012; Oz, Yang, Dinc, & Shippenberg, 2007; Spivak, Kim, Liu, Lupica, & Doyle, 2012). Here, we examine if highly lipid sensitive Kir2.1 channels, important for establishing I_K1_ in cardiac, skeletal, and smooth muscles, as well as neurons, are also regulated by endocannabinoid lipids. We have assessed the impact of various endocannabinoids from their 2 main chemical classes; fatty acid ethanolamides (FAEs) and 2-monoacylglycerols (2-MGs). Additionally, we expand our analysis to other Kir channels with intermediate sequence homology to assess if endocannabinoid regulation of Kirs is conserved across the family.

## ABBREVIATIONS

2-MGs = 2-monoacylglycerols; FAEs = fatty acid ethanolamides; AEA = Arachidonoyl Ethanolamide or anandamide; oxy-AEA = oxy-Arachidonoyl Ethanolamide; OEA = Oleoyl Ethanolamide; POEA = Palmitoleoyl Ethanolamide; LEA = Linoleoyl Ethanolamide; γ-LEA = γ-Linolenoyl Ethanolamide; α-LnEA = α-Linolenoyl Ethanolamide; DEA = Docosatetraenoyl Ethanolamide; NEA = Nervonoyl Ethanolamide; ArEA = Arachidoyl Ethanolamide; SEA = Stearoyl Ethanolamide; NAGly = Arachidonoyl Glycine; 1-AG = 1-Arachidonoyl Glycerol; 2-AG = 2-Arachidonoyl Glycerol; 2-LG = 2-Linoleoyl Glycerol; AS = N-Arachidonoyl-L-Serine; 2-PG = 2-Palmitoyl Glycerol

## MATERIALS & METHODS

### Drugs and reagents

Endocannabinoids (Cayman Chemical, USA) were pre-diluted in 99.8% ethanol at a working concentration of 10 mM. Barium Chloride (Sigma-Aldrich, USA) was diluted to a working concentration of 1 M with distilled water from a stock solution. Horse serum, penicillin-streptomycin, and kanamycin stock solutions (Thermo Fisher Scientific, Gibco Cell Culture, USA) were used pure and undiluted.

### Molecular biology and cell expression

Human Kir2.1 and Kir7.1 cDNA were previously sub-cloned into the pcDNA3.1 and pCMV6 expression vectors, respectively. Rat Kir4.1 cDNA was previously sub-cloned into the pcDNA3.1 expression vector. Briefly, linearized cDNA was obtained by linearizing Kir2.1, Kir4.1 and Kir7.1 cDNA with Mlul, Mfel and Smal (New England Biolabs), respectively. RNA was obtained by using ∼1.0 μg of linearized cDNA for an *in vitro* transcription synthesis using the mMESSAGE mMACHINE™ T7 Transcription kit (Thermo Fisher Scientific, Life Technologies, USA).

Unfertilized *Xenopus* oocytes extracted from female *Xenopus laevis* frogs were used for all electrophysiological experiments. Post-extraction, oocytes were subjected to a controlled temperature of 17 – 19 °C and placed in vials containing Barth antibiotic solution (mM): 90 NaCl, 3 KCl, 0.82 MgSO_4_.7H_2_O, 0.41 CaCl_2_.2H_2_O, 0.33 Ca(NO_3_)_2_.4H_2_O and 5 HEPES supplemented with 100 U/mL of penicillin-streptomycin and 10 mg/mL of kanamycin stock (10 mg/mL). Oocytes were microinjected with 4.6 ng of either Kir2.1, Kir4.1 and Kir7.1 RNA using a Drummond Nanoject II injector (Drummond Scientific Company). After microinjection, oocytes were incubated in Barth antibiotic solution supplemented with 5% horse serum. Cells were used for electrophysiological recordings 1 – 2 days post microinjection.

### Electrophysiological Recordings

Electrophysiological recordings were induced using the two-electrode voltage clamp (TEVC) technique. Glass borosilicate rapid fill microelectrode pipettes (FHC Inc., USA) were filled with 1 M KCl solution. Macroscopic currents from oocytes expressing Kir2.1, Kir4.1 and Kir7.1 were recorded in a bath solution containing (in mM): 89 KCl, 15 HEPES, 0.4 CaCl_2_, and 0.8 MgCl_2_, pH = 7.4 using OC-725C amplifier (Warner Instruments, USA) and digitized using a Digidata 1322A (Molecular Devices). Data was acquired with the Clampex 10.5 at a sampling rate of 5 KHz with a filter of 1 KHz. Kir2.1 inward currents were assessed by test-steps between -150 to +150 mV (ΔV = +10 mV) from a V_H_ = 0 mV, followed by a 50 ms step to 0 mV. For weaker rectifiers Kir7.1 and Kir7.4, the range of test-steps were limited from -150 to +70 mV or +100 mV respectively. Endocannabinoids were added to the oocyte containing bath in 10 µM increment after recordings stabilized from the previous condition. Ethanol was used at our vehicle for each endocannabinoid and equimolar quantities were used as controls. To ensure changes in G_max_ observed were not due to large increases in leak currents over the course of our long recordings, I-V curves were also assessed following the addition of 100 µM BaCl_2_ to the bath as the final recording of the series. Recordings were performed at room temperature.

### Data analysis and statistics

Recordings were analyzed offline using the Clampfit software (Molecular Devices, USA). The data was analyzed then plotted using the Origin 8.0 software (Northampton, MA, USA). To investigate the concentration-dependent effects of specific subsets of endocannabinoids on Kir channel currents, a pair-wise comparison methodology was utilized. Currents recorded in each cell following stabilization of currents following cannabinoid treatments were normalized to the control current (0 µM endocannabinoid) at -150 mV enabling pair-wise evaluation of the changes in currents induced by each test condition. The resulting normalized I-V curves could then be averaged, with the appropriate standard error of the mean (SEM) as presented in the figures. Each normalized I-V plot was fit with a linear relationship looking at voltages ranges from -80 to -150 mV to determine the maximal slope conductance (G_max_) at a given concentration (X µM) of the cannabinoid tested. The relative increase in the Kir function induced by each endocannabinoid (ΔG_max_) at a given concentration (X µM) was determined by evaluating:

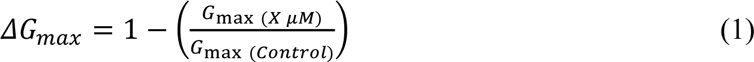

ΔG_max_ was then plotted against concentration and EC_50_ values were determined by fitting concentration dependence curves with the dose-response curve in Origin2021b (OriginLabs):

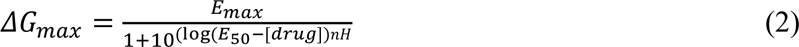

where E_max_ is the maximal effect on conductance, and nH is the hill co-efficient. Data are presented as means (±) standard error. After calculation of the EC_50_ and the E_max_ value, a 5% increase in relative G_max_ was established as a threshold to identify endocannabinoids that have an effect on Kir2.1 function based on the slight increase in current observed at high concentrations of the vehicle (ethanol) used as a solvent. Therefore, a two-sample T-test was used to test the hypothesis that E_max_ of a particular cannabinoid was greater than that of ethanol.

### Molecular Dynamics Simulations

#### System Setup

The cryo-EM structure of Kir2.1 (PDB ID: 7ZDZ) was obtained from the Protein Data Bank (Fernandes et al., 2022). The atomistic model of the protein was converted into a coarse-grained (CG) representation using martinize2 and embedded within lipid mixtures using the INSANE tool (Wassenaar, Ingolfsson, Bockmann, Tieleman, & Marrink, 2015). Elastic networks with a force constant of 500 kJ mol^-1^ nm^-2^ were employed to preserve the quaternary protein structure.

To investigate the interaction between endocannabinoids and Kir2.1, the protein was inserted into a binary mixture comprising 1-palmitoyl-2-oleoyl-sn-glycero-3-phosphocholine (POPC) and specific endocannabinoids (2-PG, 2-AG, ArEA, or AEA) within a simulation box of dimensions x=15 nm, y=15 nm, and z=15 nm.

To assess the impact of different endocannabinoids on lipid distribution and membrane properties, complex lipid bilayers were constructed using prevalent sarcolemma lipids. Simulations were conducted with and without Kir2.1, incorporating 3% of the aforementioned endocannabinoids with simulation box dimensions of x=30 nm, y=30 nm, and z=20 nm. Each simulation system was named according to the included endocannabinoid type. Detailed lipid compositions are provided in the supplementary materials, Table S1 and S2.

#### Simulation Setup

All simulations were performed using GROMACS 2021.2 (Abraham et al., 2015) simulation package with the Martini 3 force field (Souza et al., 2021). Initial systems were energy-minimized using the steepest descent algorithm for 1000 steps. Equilibration was conducted in a single step for binary mixtures and seven steps for sarcolemma systems, gradually increasing the timestep from 0.5 fs to 20 fs. Temperature was maintained at 310 K using a velocity-rescaling thermostat (Bussi, Donadio, & Parrinello, 2007) with a time constant of 1 ps, while pressure was held at 1 bar using the Berendsen barostat ps (Berendsen, Postma, Van Gunsteren, DiNola, & Haak, 1984) with a 5 ps pressure coupling time constant.

Production runs were performed using a 20 fs timestep for 80 μs (binary systems) or 60 μs (sarcolemma systems) at 310 K with the velocity-rescaling thermostat and 1 bar pressure using the Parrinello-Rahman barostat with a 12 ps pressure coupling time constant.

### Computational analyses

Density analysis for 2-PG, 2-AG, ArEA or AEA was performed using the GROMACS tool *densmap*. The last 20 μs of the binary mixtures were used for the analysis. The **depletion-enrichment index (DEI)** was calculated using an in-house protocol described previously (Corradi et al., 2018) applying a 0.8 nm cut-off distance from the embedded protein. The last 10 μs of the simulations were used for the analysis. For the **membrane thickness** analysis, FATSLiM software (Buchoux, 2017) was employed using the PO4 coarse-grained beads as headgroup selection on the last 5 μs of each trajectory. 3D visualizations were performed using the VMD (Humphrey, Dalke, & Schulten, 1996) and the PyMOL (DeLano, 2002) were used. All the figures were plotted using Python libraries such as Pandas (The pandas development team. (2020) pandas-dev/pandas: Pandas), Numpy (Harris et al., 2020), Matplotlib (Hunter, 2007) and Seaborn (Waskom, 2021).

### Protein expression and purification

Expression and purification of hKir2.1-WT were performed as described previously (Fernandes et al., 2022). Briefly, hKir2.1-WT (pPIC9K vector) was expressed in *Pichia pastoris* yeast cells (SMD1163 strain). After cell lysis with the FastPrep 24 (MP Biomedicals), membrane fractions were solubilized by adding 29.3 mM DDM (1.5% w/v, n-Dodecyl-b-D-maltoside, Glycon). Protein purification involved an affinity chromatography step using cobalt affinity resin (Takarabio), followed by size exclusion chromatography on a Superdex® 200 (10/300) GL column (Cytiva). Fractions corresponding to the tetramer were collected, flash-frozen with liquid N_2_ and stored at -80 °C.

### Surface plasmon resonance (SPR)

The interaction between hKir2.1 and two endocannabinoids, namely Arachidoyl Ethanolamide (AEA) and 2-Palmitoyl Glycerol (PAG) (Cayman Chemical), was characterized by SPR. Experiments were performed in triplicate at 25 °C on a Biacore 3000 instrument (Cytiva). Experiments were controlled by Biacore 3000 Control software v4.1, using the running buffer 20 mM Tris-HCl (pH 7.5), 150 mM KCl, 0.05 mM EDTA, and 0.03% DDM. In all experiments, a flow cell was left blank to be used as a reference for the sensorgrams (no non specific binding was observed). Activation of a carboxymethylated dextran (CM5) sensor chip was done using standard Biacore procedures, described also in (Fernandes et al., 2022; Zuniga et al., 2024). An anti-His antibody (Cytiva) at 200 nM in sodium acetate (pH 4.5) was immobilized onto the activated CM5 sensor chip (flow 10 μL/min, contact time 7 min), and saturation was achieved with 1 M ethanolamine HCl (pH 8.5). hKir2.1-WT at 50 nM, diluted in running buffer, was then immobilized on the anti-His antibody (flow 5 μL/min, contact time 5 min). Binding of the two endocannabinoids was assessed by single injections of AEA and PAG at 10 µM and 50 µM diluted in running buffer (flow 20 μL/min, contact time 3 min) over both the reference cell and the ligand cell.

## RESULTS

### Endocannabinoids enhance Kir2.1 function

We examined the effects of endocannabinoids on Kir2.1 function by expressing these channels in *Xenopus* oocytes, because they lack cannabinoid receptors (CBRs) (Karimi et al., 2018). Endocannabinoids generally fall into two chemical classes based on their headgroups (Fatty acid ethanolamines (FAEs) and 2-monoacylglycerols (2-MGs)). In some cases, the glycerol group can be substituted with other moieties such as serine (as in arachidonoyl serine (AS)) or glycine (as in N-arachidonoyl glycine (NAGly)). Kir2.1 currents were recorded every 5 mins until the I-V curves were stable. Once stable, individual endocannabinoids were added to the bath in 10 µM increments only after currents stabilized at the previous concentration (typically between 20 – 60 mins). To enable cell-to-cell comparisons of the effects of each cannabinoid, each I-V recorded on a single cell was normalized to the stabilized control currents measured at -150 mV in 0 µM endocannabinoid. This enabled averaging across cells for a given endocannabinoid treatment and assessment of concentration dependencies. Since ethanol was used as the solvent to dissolve all endocannabinoids, we performed similar experiments with the equivalent quantities of ethanol for each concentration of endocannabinoids tested. We observe only a 3.5 ± 0.7% increase in Kir2.1 currents due to ethanol (Fig. 1B, 2A, and Table 1).

**Figure 1.**
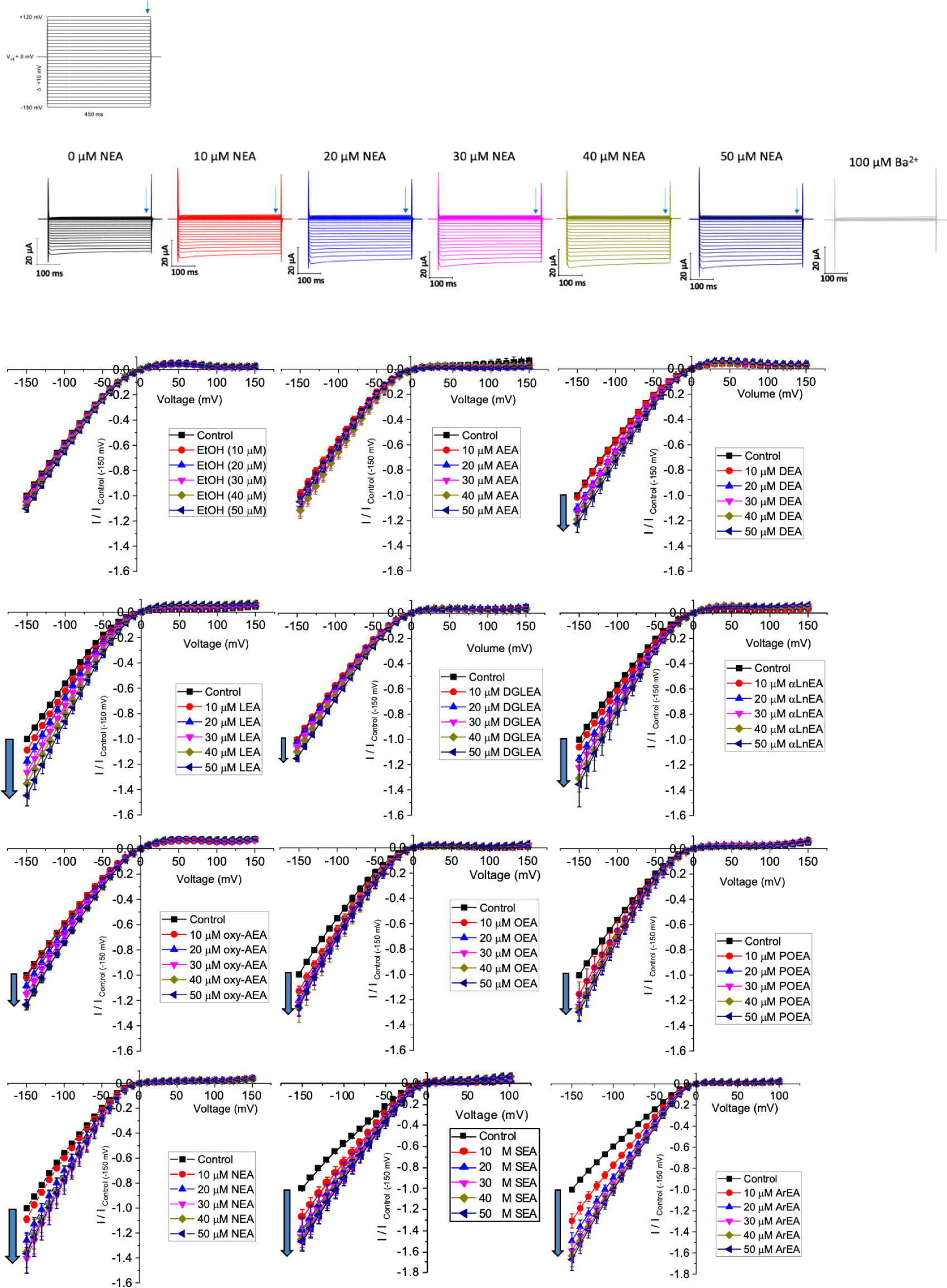
Effect of fatty acid ethanolamides (FAEs) on Kir2.1 currents. Current-voltage (I-V) relationship of Kir2.1 channels expressed in Xenopus laevis oocytes following the incremental addition of the listed endocannabinoids. For each cell, currents were normalized to the current elicited at -150 mV for control conditions (0 µM endocannabinoid). This enabled the averaging of pair-wise data from different cells.

**Figure 2.**
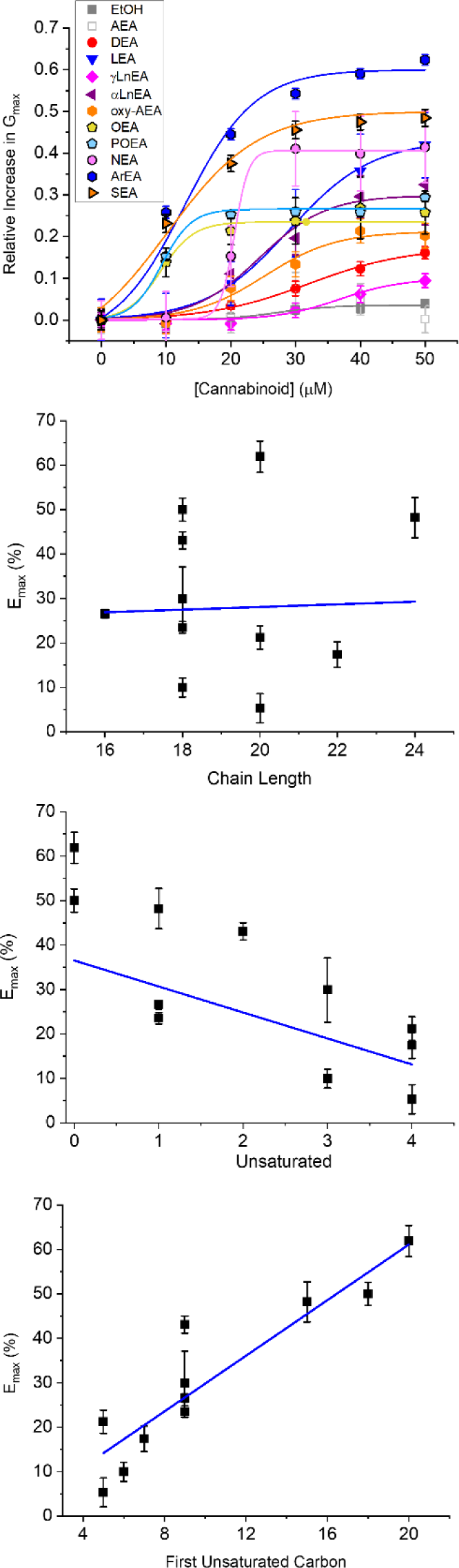
Concentration dependence of FAEs on Kir2.1 current. **(A)** Concentration dependence curves were calculated for each endocannabinoid by plotting the relative increase in slope conductance (Gmax) compared to control. Data was then fit to a dose-response function to determine EC_50_ and maximal effect (E_max_). **(B)** There is no correlation between Emax and FAE endocannabinoid chain-length. However, there is a co-relation between Emax and the number of unsaturated bonds **(C)** (P = 0.05) and the position of the first unsaturated carbon **(D)** (P = 0.0008).

**Table 1.** Maximal effect (E_max_) and the EC_50_ for each fatty acid ethanolamine (FAE) endocannabinoid tested. A 5% increase in current was established as a threshold to identify endocannabinoids that have an effect on Kir2.1 function based on the slight increase in current observed at high concentrations of the vehicle (Ethanol) used as a solvent.

| **Cannabinoid** | **$EC_{50}$ (μM)** | **$E_{\max}$ (%)** |
| --- | --- | --- |
| **Ethanol (Vehicle)** | | $3.5 \pm 0.7$ |
| **AEA** | | $5.3 \pm 1.3$ |
| **DEA** | $32.0 \pm 3.9$ | $17.4 \pm 2.8$ * |
| **LEA** | $29.0 \pm 1.0$ | $43.1 \pm 1.9$ * |
| **YLnEA** | $36.8 \pm 3.8$ | $10.0 \pm 2.1$ * |
| **αLnEA** | $24.5 \pm 4.2$ | $29.9 \pm 7.2$ * |
| **oxy-AEA** | $25.3 \pm 1.1$ | $21.2 \pm 2.6$ * |
| **OEA** | $9.2 \pm 0.9$ | $23.5 \pm 1.3$ * |
| **POEA** | $9.3 \pm 0.7$ | $21.2 \pm 1.0$ * |
| **NEA** | $23.3 \pm 2.1$ | $48.2 \pm 4.5$ * |
| **ArEA** | $12.6 \pm 4.1$ | $61.9 \pm 3.5$ * |
| **SEA** | $10.2 \pm 4.9$ | $50.0 \pm 0.1$ * |

We first examined the effect of 11 FAEs on Kir2.1 function, by pair-wise examination of their effect on maximal slope conductance (G_max_) (Figs. 1 and 2). Notably, the best studied FAE endocannabinoid anandamide (AEA) had no effect on Kir2.1 currents. However, we observe differing effects of the remaining 10 FAEs on Kir2.1 currents (Table 1) with changes in maximal effect (E_max_) ranging from approximately 15 – 60% increase in G_max_. ArEA has the largest effect on Kir2.1, increasing G_max_ by 61.9 ± 3.5%, while OEA, POEA, ArEA and SEA are the most potent FAEs tested, with EC_50_ approximately 10 µM.

To determine if a specific physiochemical property of the FAEs is the molecular driver for stimulating Kir2.1 currents, we assessed the correlation between the E_max_ for each FAE and the chain length (Fig. 2B), degree of unsaturation of the acyl tail (Fig. 2C) and the position of the first unsaturated carbon (Fig. 2D). Notably, there is no correlation between FAE chain length and the effect on Kir2.1 currents. However, there is strong correlation between Kir2.1 function and both the position of the first unsaturated carbon and degree of unsaturation.

We then expanded our examination of endocannabinoids to include 2-MGs (Fig. 3). Endocannabinoids in this class contains a glycerol head group at the sn-2 position rather than an ethanolamide head group, followed by fatty acid chains of varying lengths and unsaturation. Like what we observe for the FAE class, 2-MG endocannabinoids had varying effects on Kir2.1 function. Notably, 2-PG, NAGly and 2-AG produce the largest increases to Kir2.1 currents with E_max_ values of 50.8 ± 2.7%, 33.8 ± 3.8% and 22.4 ± 0.9 %, respectively, while 2-LG and AS have no discernable effects (Fig. 3 and Table 2). It is not possible for us to assess the correlation between chemical properties and Kir2.1 function for this class because there are fewer commercially available 2-MG endocannabinoids, and the range of their physiochemical diversity is more limited than the FAEs available.

**Figure 3.**
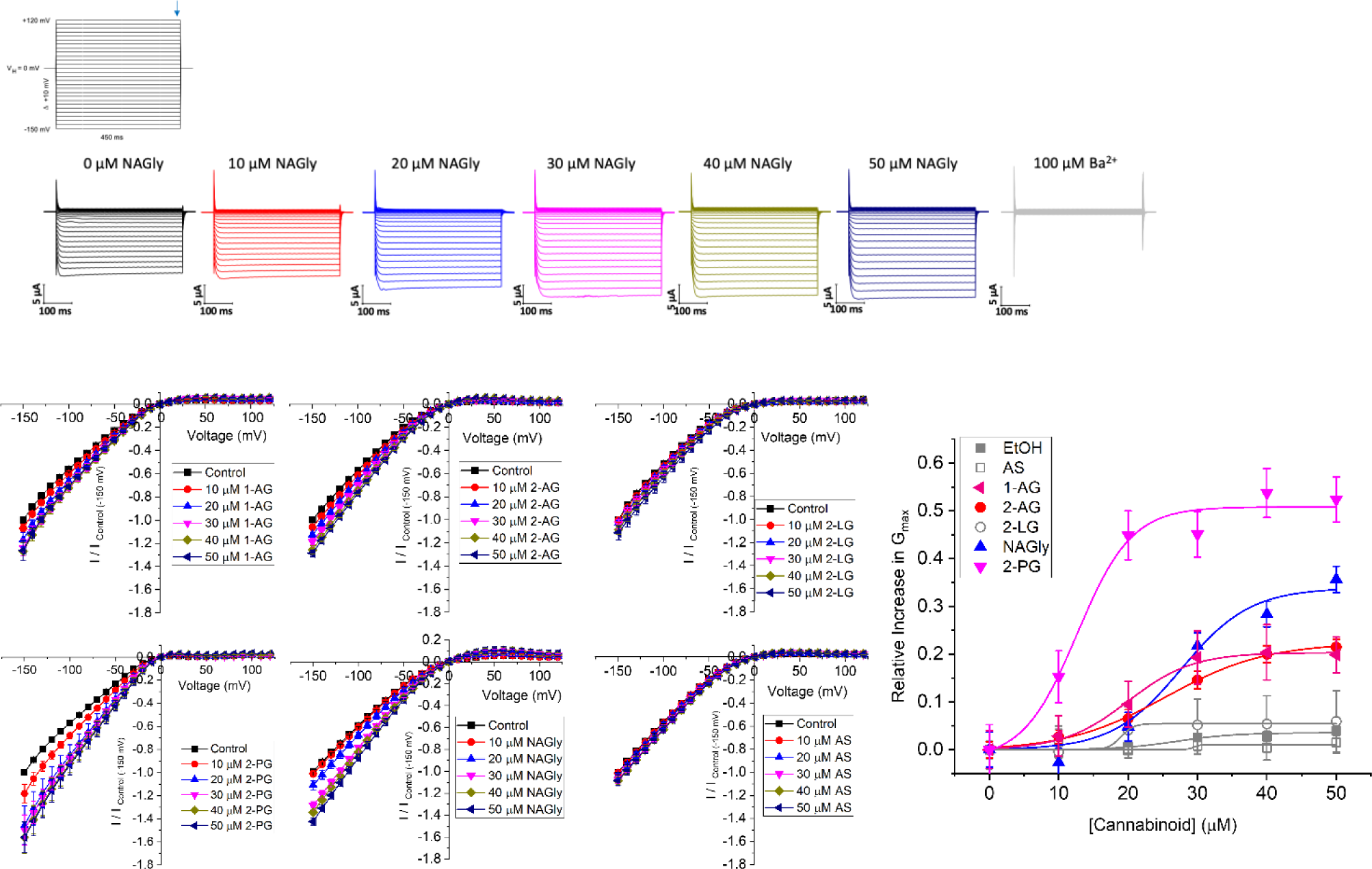
Concentration dependence of 2-MGs on Kir2.1 current. (A) Current-voltage (I-V) relationship of Kir2.1 channels following the incremental addition of the listed endocannabinoids. For each cell, currents were normalized to the current elicited at -150 mV for control conditions (0 µM endocannabinoid). This enabled the averaging of pair-wise data from different cells. (B) Concentration dependence curves were calculated for each endocannabinoid by plotting the relative increase in slope conductance (Gmax) compared to control. Data was then fit to a dose-response function to determine EC_50_ and maximal effect (E_max_).

**Table 2.** Maximal effect (E_max_) and the EC_50_ for each 2-monoacylglycerol (2-MG) and related endocannabinoid tested. A 5% increase in current was established as a threshold to identify endocannabinoids that have an effect on Kir2.1 function based on the slight increase in current observed at high concentrations of the vehicle (Ethanol) used as a solvent.

| **Cannabinoid** | **$EC_{50}$ (μM)** | **$E_{\max}$ (%)** |
| --- | --- | --- |
| **Ethanol (Vehicle)** | | $3.5 \pm 0.7$ |
| **AS** | | $1.0 \pm 0.1$ |
| **1-AG** | $20.0 \pm 0.9$ | $20.3 \pm 0.6$ * |
| **2-AG** | $25.5 \pm 1.1$ | $22.4 \pm 0.9$ * |
| **2-LG** | | $5.4 \pm 0.3$ |
| **NAGly** | $27.7 \pm 2.8$ | $33.8 \pm 3.8$ * |
| **2-PG** | $12.7 \pm 2.0$ | $50.8 \pm 2.7$ * |

### Assessing the conservation of endocannabinoid effects on other Kir channels

To determine if the endocannabinoid regulation of the strong rectifier Kir2.1 is conserved across the inward rectifier family, we examined the effects of the several effective endocannabinoids on the intermediate rectifying Kir7.1 (Fig. 4) and weak rectifying Kir4.1 channels (Fig. 5 and Table 3). OEA, LEA, ArEA, and 2-PG had no observable effect on Kir7.1 channels that differed from vehicle. On the other hand, Kir4.1 channels appear more sensitive to modulation by endocannabinoids than Kir2.1 channels. ArEA induces a 144 ± 23.8% increase in G_max_ with an EC_50_ **=** 4.5 ± 3.9 µM, while 2-PG increases Kir4.1 G_max_ by 164 ± 7.9% with an EC_50_ = 12.4 ± 0.7 µM. These data indicate that the mechanism(s) of endocannabinoid regulation of Kir channels is not conserved across the family.

**Figure 4.**
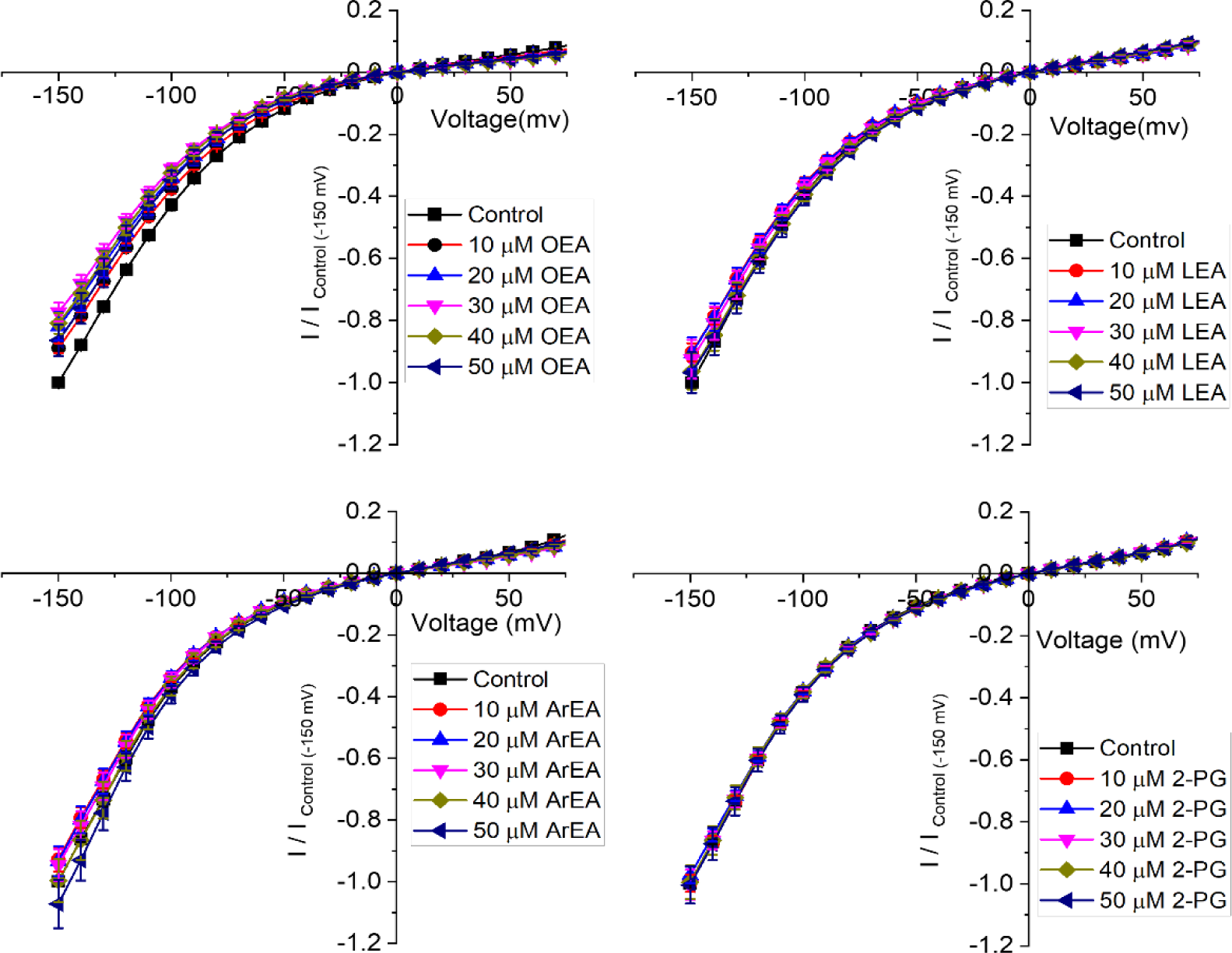
Endocannabinoids do not affect Kir7.1 currents. **(A-D)** Current-voltage (I-V) relationship of Kir7.1 channels following the incremental addition of the listed endocannabinoids. For each cell, currents were normalized to the current elicited at -150 mV for control conditions (0 µM endocannabinoid). This enabled the averaging of pair-wise data from different cells. Endocannabinoids that had a large effect on Kir2.1 currents did not alter Kir7.1 currents over the same time-course of application.

**Figure 5.**
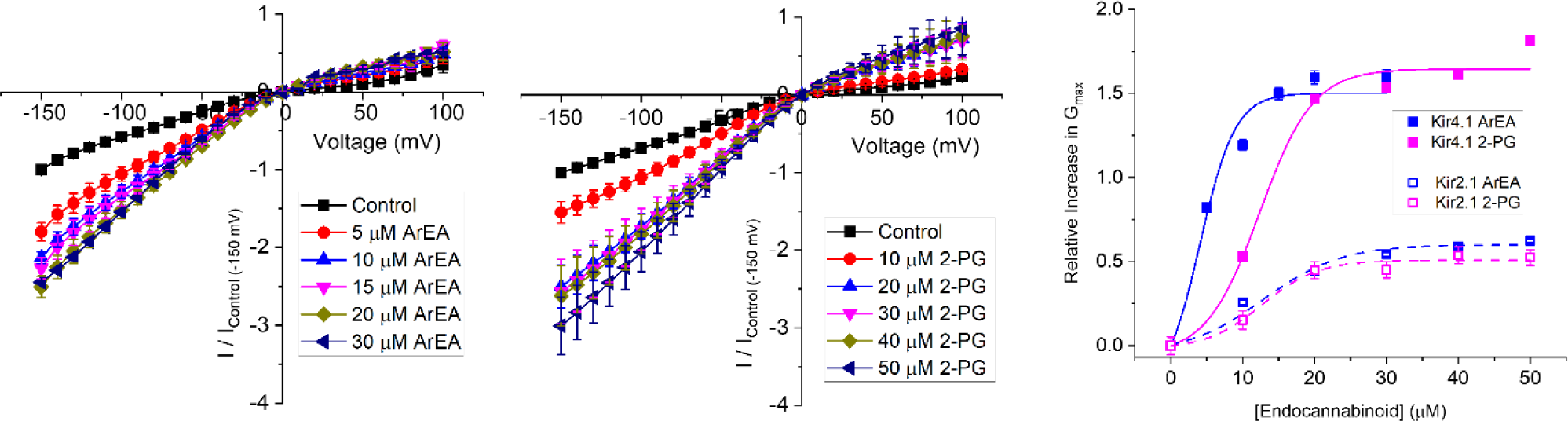
Endocannabinoids have a strong effect on Kir4.1 currents. Current-voltage (I-V) relationship of Kir4.1 channels following the incremental addition of the **(A)** ArEA or **(B)** 2-PG. For each cell, currents were normalized to the current elicited at -150 mV for control conditions (0 µM endocannabinoid). This enabled the averaging of pair-wise data from different cells. **(C)** Concentration dependence curves were calculated for each endocannabinoid by plotting the relative increase in slope conductance (G_max_) compared to control. Data was then fit to a dose-response function to determine EC_50_ and maximal effect (E_max_). For comparison, data from Kir2.1 channels are shown in open squares with dashed lines.

**Table 3.** Comparison of the maximal effect (E_max_) and the EC_50_ for ArEA and 2-PG on Kir2.1 and Kir4.1 channels.

| **Cannabinoid** | **$EC_{50}$ (μM)** | **$E_{\max}$ (%)** |
| --- | --- | --- |
| **Kir2.1 ArEA** | $12.6 \pm 4.1$ | $61.9 \pm 3.5$ |
| **Kir2.1 2-PG** | $12.7 \pm 2.0$ | $50.8 \pm 2.7$ |
| **Kir4.1 ArEA** | $4.5 \pm 3.9$ | $144 \pm 23.8$ |
| **Kir4.1 2-PG** | $12.4 \pm 0.7$ | $164 \pm 7.9$ |

### Endocannabinoids do not directly interact with Kir2.1 channels

To assess the potential interactions between the endocannabinoids and Kir2.1, we modelled different FAEs and 2-MGs using the Martini 3 building blocks (Souza et al., 2021). These models were embedded within a lipid mixture composed of 1-palmitoyl-2-oleoyl-sn-glycero-3-phosphocholine (POPC) and the respective endocannabinoids. MD simulations, spanning 80 μs, were conducted to observe the spatial distribution of endocannabinoids around Kir2.1. The 2D density maps generated from these simulations revealed that representatives of the 2-MG family and the FAE family did not exhibit significant accumulation near Kir2.1 (Fig. 6A). Although slightly higher densities of anandamide (AEA) were observed in proximity to Kir2.1, further analysis via residence time did not indicate sustained interactions with the channel.

**Figure 6.**
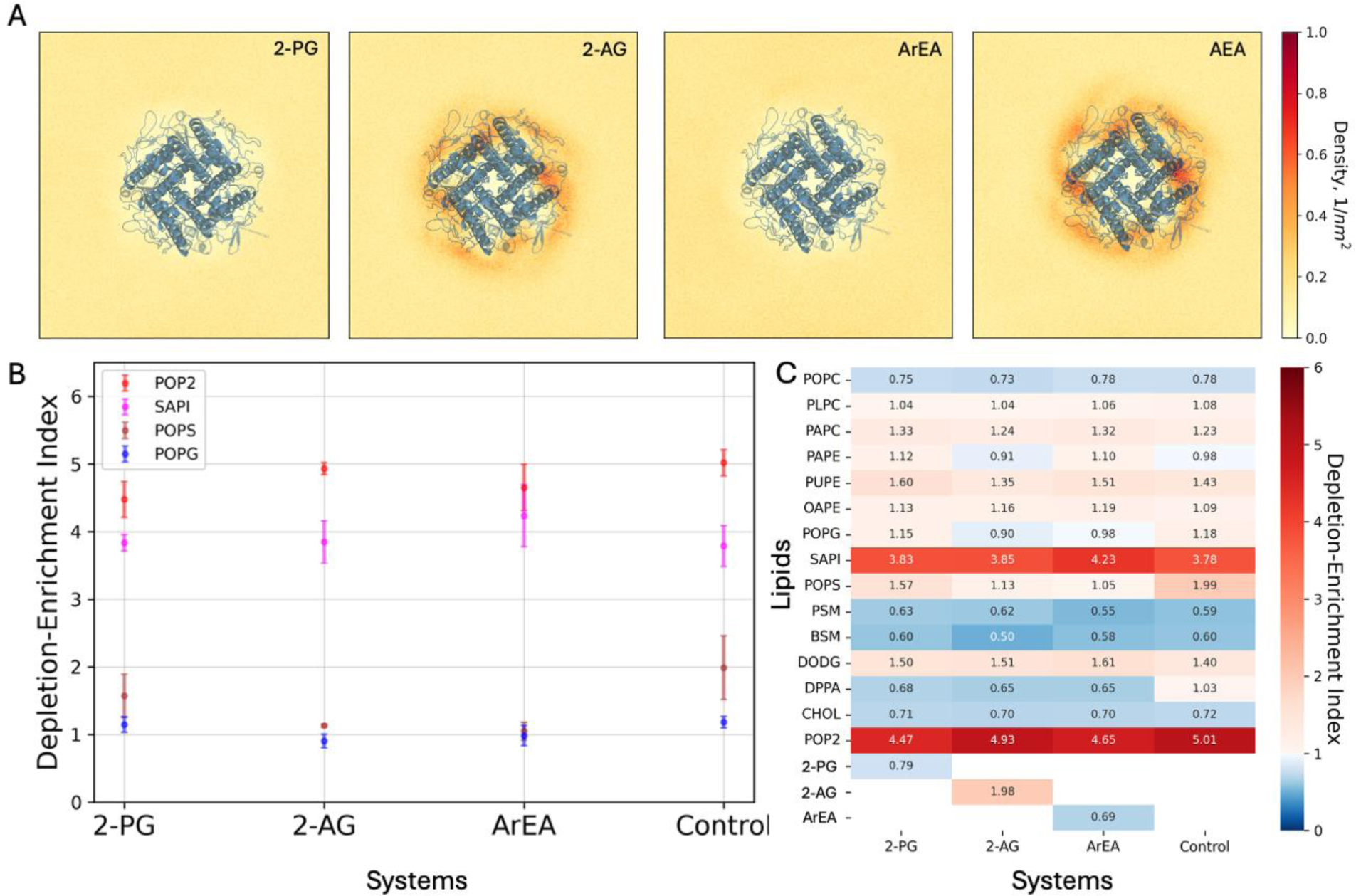
Coarse-grained simulations of Kir2.1 channels in the presence of endocannabinoids. (A) 2D density maps computed from the final 20 μs of the binary mixtures MD simulations depicting the spatial distribution of the endocannabinoids around the Kir2.1. (B) Lipid Depletion-Enrichment Index (DEI) calculated for negatively charged lipids in sarcolemma models, employing a 0.8 nm cutoff distance from Kir2.1. The values represent averages and standard errors derived from three independent replicates. (C) Average DEI values across three replicates for each lipid within the system, providing insights into overall membrane lipid redistribution.

To explore the potential indirect effects of endocannabinoids on Kir2.1 conductance, we performed extensive MD simulations (∼ 60 μs per replica) on complex sarcolemma models with and without Kir2.1 embedded. These models include prevalent lipids found in the cardiac sarcolemma and either 2-PG, 2-AG, or ArEA modelled at the previous step. Control simulations without endocannabinoids were also performed for comparison.

Analysis of the depletion-enrichment index (DEI) for all lipids within the system, including endocannabinoids, revealed distinct effects on membrane lipid redistribution. Notably, phosphatidylinositol 4,5-bisphosphate (PIP2) exhibited enrichment around Kir2.1 across all systems, albeit with varying DEI values. Particularly, the system containing 2-PG displayed the lowest DEI value compared to control and other endocannabinoid systems (Fig. 6C). The computed average DEI values with the standard errors for the negatively charged lipids also represent the difference in lipid enrichment or depletion (Fig. 6B).

Additionally, we analyzed membrane thickness in the sarcolemma models during the final 5 μs of simulations. The systems containing Kir2.1 exhibited similar trends in membrane thickness compared to those without Kir2.1. Specifically, the systems with ArEA and Kir2.1 demonstrated the highest thickness, followed by 2-PG system, while the control systems devoid of endocannabinoids exhibited the lowest thickness (Table 4).

**Table 4.** Thickness analysis of the complex sarcolemma membrane models with and without Kir2.1 embedded. Thickness measurements were obtained during the final 5 μs of simulations using PO4 beads for lipid headgroup selection. Average values and standard errors were calculated across three replicates.

|  | **Systems** | **Thickness, Å** |
| --- | --- | --- |
| **Lipid-only systems** | 2-PG | 41.163 +- 0.002 |
|  | 2AG | 41.084 +- 0.002 |
|  | ArEA | 41.274 +- 0.002 |
|  | Control | 40.992 +- 0.002 |
| **Kir2.1 systems** | 2-PG | 41.139 +- 0.002 |
|  | 2AG | 41.063 +- 0.002 |
|  | ArEA | 41.246 +- 0.002 |
|  | Control | 40.958 +- 0.002 |

Our computational analyses suggest that endocannabinoids, including 2-PG and ArEA, do not directly interact with Kir2.1 channels. However, their presence modulates membrane lipid redistribution, potentially influencing Kir2.1 function indirectly. These results were further confirmed by assessing the binding efficiency between Kir2.1 and two endocannabinoids (ArEA, 2-PG) by surface plasmon resonance (Fig. 7). His-tagged Kir2.1 was fixated at 1830 RU (response units) onto an immobilized anti-His antibody (at 10200 RU) on a CM5 sensor chip. Single injections of ArEA and 2-PG at 10 μM and 50 μM revealed no binding with Kir2.1, highlighting an indirect effect of the endocannabinoids on the functional properties of Kir2.1.

**Figure 7.**
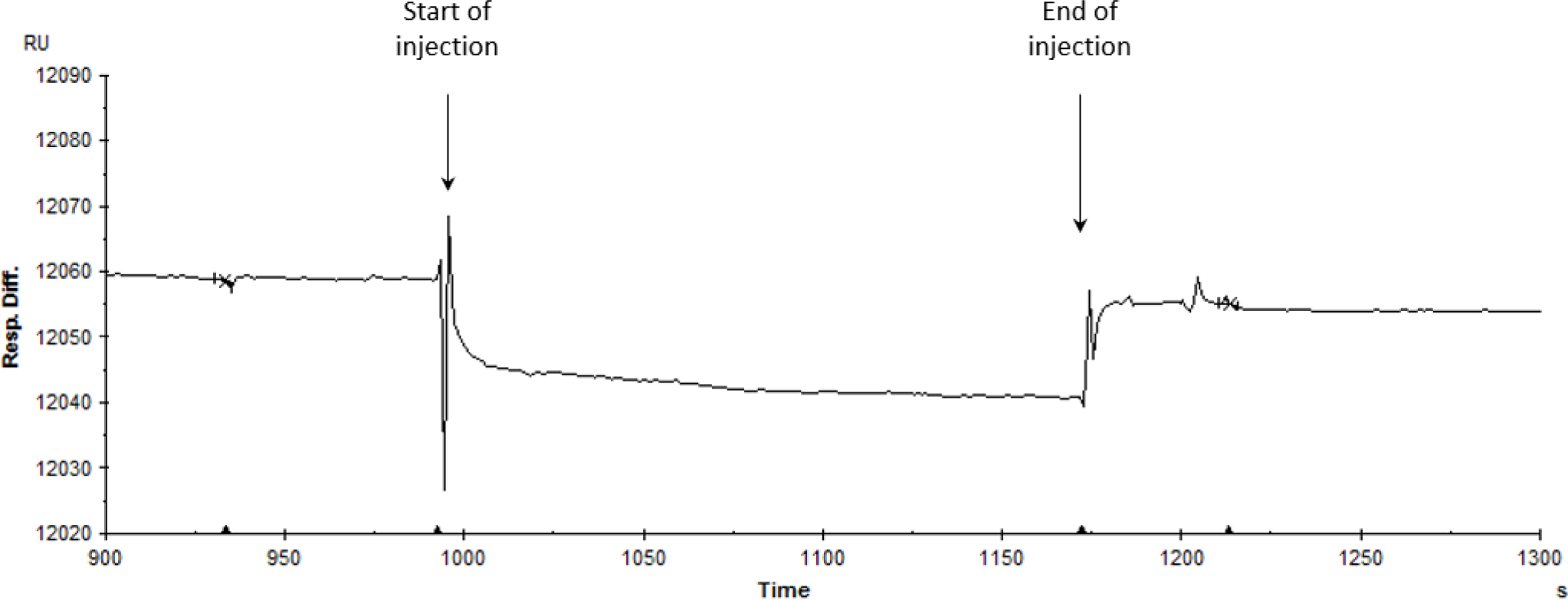
Assessment of the interaction between hKir2.1-WT and endocannabinoids. Single injection of ArEA at 50 μM on a His-tagged hKir2.1 captured by an anti-His antibody covalently immobilized-CM5 sensor chip (at 60 μL/3 min). The SPR response is expressed in response units relative to the time in seconds. The decrease in response difference (20 relative units RU) corresponds to the buffer effect and is indicative of no interaction.

### Endocannabinoid effects on LQT7 mutant Kir2.1 channels

Since endocannabinoids increase Kir2.1 conductance, we considered as proof of concept if enriching endocannabinoids could be useful in enhancing Kir2.1 mutant channels linked to LQT7 or Andersen-Tawil Syndrome. As a test example, we worked with G144S mutant Kir2.1. Since G144S Kir2.1 alone does not express any functional current, we co-expressed with equivalent amount of WT Kir2.1 to mimic heterozygous patients, which have been shown to express ∼40% of WT Kir2.1 currents (Tristani-Firouzi et al., 2002). We demonstrate that endocannabinoids ArEA and 2-PG still have an enhancing effect on WT:G144S Kir2.1 currents, however, the concentration dependencies differ from WT alone (Fig. 8). Specifically, the LQT7 mutation reduces the efficacy and/or potency of these endocannabinoids. ArEA induces a maximal change in conductance (E_max_) of 39.0 ± 6.0% in WT:G144S channels, with an EC_50_ of 30.7 ± 3.4 µM. The E_max_ induced by 2-PG is 33.1 ± 1.2% with an EC_50_ of 11.7 ± 1.0 µM. Thus, while disease-linked mutations of Kir2.1 channels can still be modulated by endocannabinoids, the effectiveness of any approach needs to be carefully (and perhaps individually) considered.

**Figure 8.**
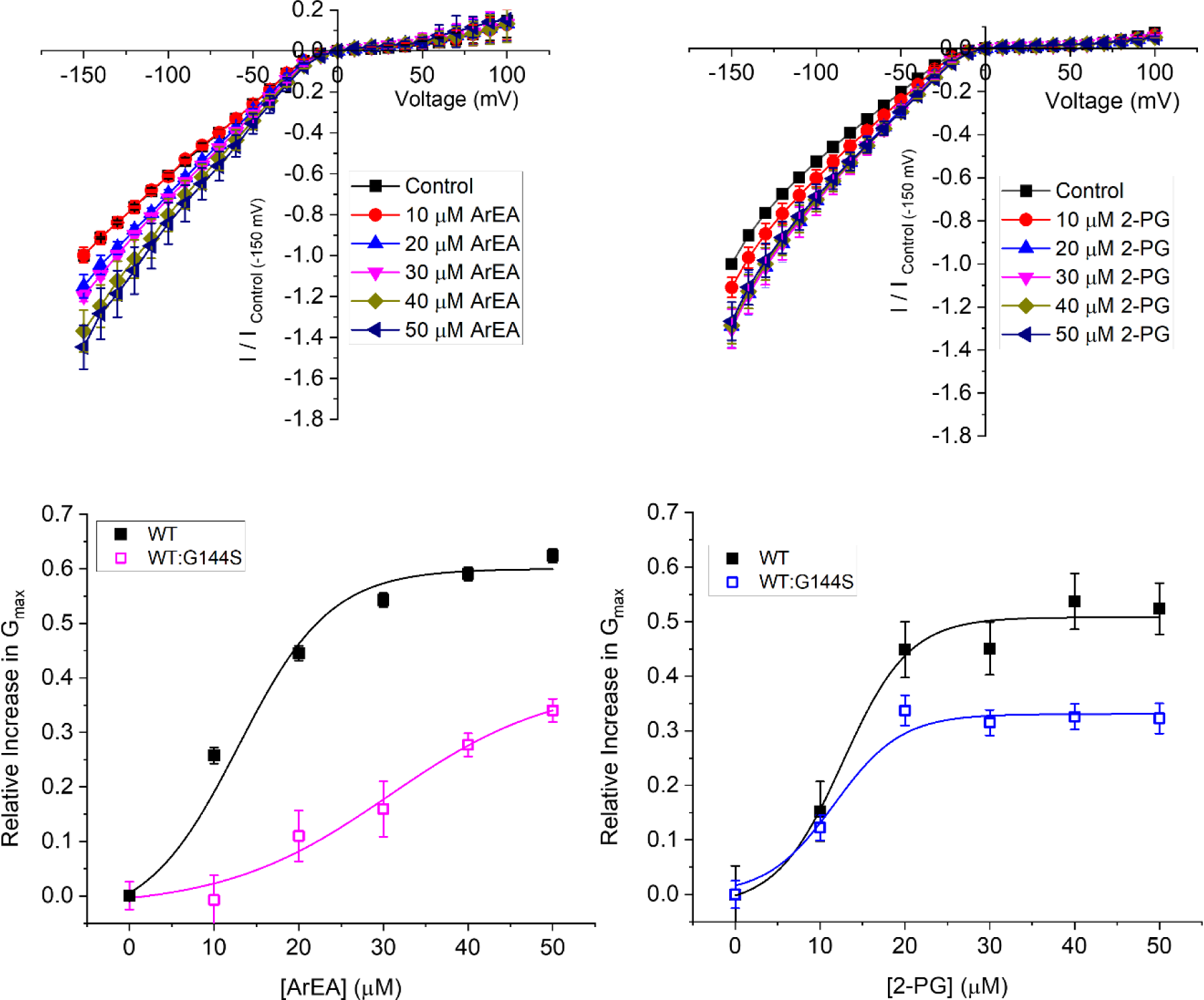
Endocannabinoids alter mutant Kir2.1 channels linked to Andersen-Tawil Syndrom / LQT7. Current-voltage (I-V) relationship of **(A)** WT Kir2.1 channels (black squares) or **(B)** 1:1 WT:G144S Kir2.1 channels following the incremental addition of the ArEA or 2-PG. For each cell, currents were normalized to the current elicited at -150 mV for control conditions (0 µM endocannabinoid). This enabled the averaging of pair-wise data from different cells. **(C & D)** Concentration dependence curves were calculated for each endocannabinoid by plotting the relative increase in slope conductance (Gmax) compared to control. Data was then fit to a dose-response function to determine EC_50_ and maximal effect (E_max_).

## DISCUSSION

A growing number of ion channels have been shown to be regulated by specific components of lipid membranes (Bolotina, Omelyanenko, Heyes, Ryan, & Bregestovski, 1989; Bowles, Heaps, Turk, Maddali, & Price, 2004; W. W. L. Cheng et al., 2011; D’Avanzo et al., 2010; Enkvetchakul et al., 2005; Furst et al., 2014; Heaps, Tharp, & Bowles, 2005; Mayar et al., 2022; Romanenko, Rothblat, & Levitan, 2002; Rosenhouse-Dantsker et al., 2013)). Endocannabinoids and endocannabinoid-like lipids also regulate the function of a variety of channels independent of cannabinoid receptor signaling mechanisms family (Ahrens et al., 2009; De Petrocellis & Di Marzo, 2011; De Petrocellis et al., 2012; Ghovanloo et al., 2021; Ghovanloo et al., 2018; Hejazi et al., 2006; Iannotti, Hill, et al., 2014; Iannotti, Silvestri, et al., 2014; Mayar et al., 2022; Okada et al., 2005; Page & Ruben, 2024; Ross et al., 2008).

Here we assess the effect of endocannabinoids in both the FAE and 2-MG classes on Kir channels expressed in *Xenopus laevis* oocytes. Unfertilized *Xenopus oocytes* lack CBRs (Xenbase.org) (Karimi et al., 2018) and thus, provide an ideal system to examine the direct effects of endocannabinoids on ion channel function in the absence of cannabinoid receptors. Our data demonstrates that in the FAE class of endocannabinoids, ArEA has the largest effect (measured by increasing G_max_), while OEA, POEA, ArEA and SEA are the most potent of those tested, with EC_50_ near 10 µM. Intriguingly, the best studied endocannabinoid anandamide (AEA) had no effect on Kir2.1 currents. Of the 2-MGs studied, 2-PG, NAGly, 1-AG and 2-AG produce the largest increases to Kir2.1 currents, with 2-PG being the most potent.

Since lipids, including endocannabinoids, can regulate the function of channels, receptors, and transporters, by direct protein-lipid interactions, or through changes in the physiochemical properties of the bilayer, we assessed the potential mechanism of action via computational and biochemical experiments. The 2D density maps generated from coarse-grained simulations revealed that endocannabinoids did not significantly accumulate near Kir2.1 (Fig. 6A). This was also supported by SPR experiments, which also failed to provide evidence to support specific binding of endocannabinoids to purified Kir2.1 channels. This suggests that endocannabinoids regulate Kir channels via indirect changes in membrane properties, rather than through direction protein-lipid interactions. These indirect effects may involve changes in membrane thickness, as suggested from our coarse-grained simulations (Table 4).

Inward rectifier channels have a highly conserved topology (Fernandes et al., 2022; Kuo et al., 2003; Tao, Avalos, Chen, & MacKinnon, 2009; Whorton & MacKinnon, 2011; Zangerl-Plessl et al., 2020), despite sequence and functional diversity in key regions. For example, Kir3 channels can bind Na^+^ and Gβγ subunits (Whorton & MacKinnon, 2011), and Kir6 members can bind ATP (Sung et al., 2022). Similarly, Kir channels have different sensitivities to phosphoinositides (Rohacs, Chen, Prestwich, & Logothetis, 1999; Rohacs et al., 2003) and cholesterol (Romanenko et al., 2004; Rosenhouse-Dantsker, Leal-Pinto, Logothetis, & Levitan, 2010). Thus, we set out to determine if endocannabinoid regulation of the strong rectifier Kir2.1 is conserved across the inward rectifier family. Notably, we observed that ArEA and 2-PG had no effect on the intermediate rectifying Kir7.1 (Fig. 4), despite the large changes they induce in Kir2.1 function. On the other hand, Kir4.1 channels, which are intermediate rectifiers, are more sensitive to ArEA compared to Kir2.1 channels (with an EC_50_ of 4.5 µM compared to 12.6 µM). Moreover, both ArEA and 2-PG induce larger changes in Kir4.1 conductance (E_max_) than those observed for Kir2.1. Thus, endocannabinoid regulation of Kir channels is not conserved among family members. Studies of 2-AG and AEA in K_ATP_ channels provide further support that endocannabinoid regulation is not conserved within the Kir family. 2-AG inhibited mouse insulinoma R7T1 β-cells with an IC_50_ of 1 µM (Spivak et al., 2012). Furthermore, K_ATP_ expressed in *Xenopus* oocytes were inhibited by AEA (CBR independent) with an IC_50_ of 8.1 µM (Oz et al., 2007), but activated in rat ventricular myoctyes in a CB2 dependent manner (Li et al., 2012). Notably, the EC_50_’s of the endocannabinoids studied here on Kir2.1 and Kir4.1 are in a similar range of IC_50_/EC_50_’s identified for a variety of other channels (Lin, 2021), and thus physiologically relevant.

Mutations in Kir2.1 channels that lead to a reduction in I_K1_ are linked to Long QT 7 (LQT7) or Andersen-Tawil Syndrome (ATS). Since endocannabinoids can increase Kir2.1 function, we assessed whether an LQT7/ATS linked mutation can be used as a rescue strategy, as a proof-of-concept. This has potential to be an effective strategy because numerous endocannabinoid-like lipids (including 2-PG, 2-LG, OEA, POEA, SEA and others) have been shown to have little or no affinity for CB1 or CB2 receptors (see (Rahman, Uyama, Hussain, & Ueda, 2021) for overview). Since homozygous mutations are entirely non-functional, we examined 2-PG and ArEA effects on 1:1 WT:G144S expressed channels, to mimic heterozygous patients, which have been shown to express ∼40% of WT Kir2.1 currents. We demonstrate that ArEA and 2-PG still have an enhancing effect on WT:G144S Kir2.1 currents, however, the concentration dependencies differ from WT alone. Specifically, the LQT7 mutation reduces the efficacy and/or potency of these endocannabinoids, depending on the lipid. This presents an important caveat to future considerations of endocannabinoid treatment of disease linked mutations. While disease-linked mutations of Kir2.1 channels, or any other channel, may be modulated by endocannabinoids or endocannabinoid-like lipids, the effectiveness of any approach needs to be carefully (and perhaps individually) considered.

Overall, we demonstrate a novel mechanism of regulation of the Kir channel family. Endocannabinoids and endocannabinoid-like lipids can alter the conductance of Kir function, however, the affinity and maximal effects are not conserved among members of the family. These effects appear to be driven by changes in the membrane properties, rather than via direct interactions of the endocannabinoid with the Kir channel. Lastly, endocannabinoids may provide an effective therapeutic approach to current rescue, however, differences in potency or maximal effects may differ from WT channels, and should be considered.

## ACKNOWLEDGEMENTS

MB wants to thank Haydee Mesa-Galloso for providing the scripts for DEI calculations. This work was supported by a Discovery Grant (RGPIN-2019-00373) from the National Science and Engineering Research Council (NSERC) and Project Grants from the Canadian Institutes of Health Research (CIHR) (FRN 173388 awarded to ND and PJT-180245 awarded to DPT). This work was also supported by AFM-Téléthon #23207 for C.V.-B, AZ and TB, and the Canada Research Chairs program (DPT). Calculations were carried out on Digital Alliance of Canada resources, supported by the Canada Foundation for Innovation and partners. SM, AA, MML, MM-Y were kindly supported by scholarships awarded by the Université de Montréal.

