## Supplementary Data for "Endocannabinoid regulation of inward rectifier potassium (Kir) channels"

**Table S1**. An example of the binary system configurations.

| Number of lipids | Lipids | | Systems | | | |
| --- | --- | --- | --- | --- | --- | --- |
|  |  |  | 2-PG | 2-AG | ArEA | AEA |
|  | Upper  leaflet | POPC | 291 | 291 | 291 | 291 |
|  |  | Endocannabinoids | 32 | 32 | 32 | 32 |
|  | Lower  leaflet | POPC | 293 | 293 | 293 | 293 |
|  |  | Endocannabinoids | 32 | 32 | 32 | 32 |

**Table S2.** An example of the sarcolemma membrane with Kir2.1 embedded system configurations.

| Number of lipids | Lipids | | Systems with Kir2.1 | | | |
| --- | --- | --- | --- | --- | --- | --- |
|  |  |  | **2-PG** | **2-AG** | **ArEA** | **Control** |
|  | Upper  leaflet | POPC | 284 | 284 | 284 | 300 |
|  |  | PLPC | 203 | 203 | 203 | 220 |
|  |  | PAPC | 257 | 257 | 257 | 272 |
|  |  | PAPE | 3 | 3 | 3 | 3 |
|  |  | PUPE | 7 | 7 | 7 | 7 |
|  |  | OAPE | 3 | 3 | 3 | 3 |
|  |  | POPG | 0 | 0 | 0 | 0 |
|  |  | SAPI | 0 | 0 | 0 | 0 |
|  |  | POPS | 0 | 0 | 0 | 0 |
|  |  | DPPA | 0 | 0 | 0 | 0 |
|  |  | PSM | 106 | 106 | 106 | 105 |
|  |  | BSM | 43 | 43 | 43 | 43 |
|  |  | DODG | 25 | 25 | 25 | 25 |
|  |  | CHOL | 505 | 505 | 505 | 501 |
|  |  | POP2 | 0 | 0 | 0 | 0 |
|  |  | Endocannabinoids | 41 | 41 | 41 | 0 |
|  | Lower  leaflet | POPC | 175 | 175 | 175 | 188 |
|  |  | PLPC | 125 | 125 | 125 | 138 |
|  |  | PAPC | 158 | 158 | 158 | 171 |
|  |  | PAPE | 52 | 52 | 52 | 52 |
|  |  | PUPE | 133 | 133 | 133 | 134 |
|  |  | OAPE | 53 | 53 | 53 | 54 |
|  |  | POPG | 15 | 15 | 15 | 16 |
|  |  | SAPI | 51 | 51 | 51 | 51 |
|  |  | POPS | 15 | 15 | 15 | 16 |
|  |  | DPPA | 15 | 15 | 15 | 16 |
|  |  | PSM | 6 | 6 | 6 | 6 |
|  |  | BSM | 2 | 2 | 2 | 2 |
|  |  | DODG | 29 | 29 | 29 | 29 |
|  |  | CHOL | 576 | 576 | 576 | 581 |
|  |  | POP2 | 23 | 23 | 23 | 24 |
|  |  | Endocannabinoids | 47 | 47 | 47 | 0 |

**Table S3.** The modelled endocannabinoid structures.

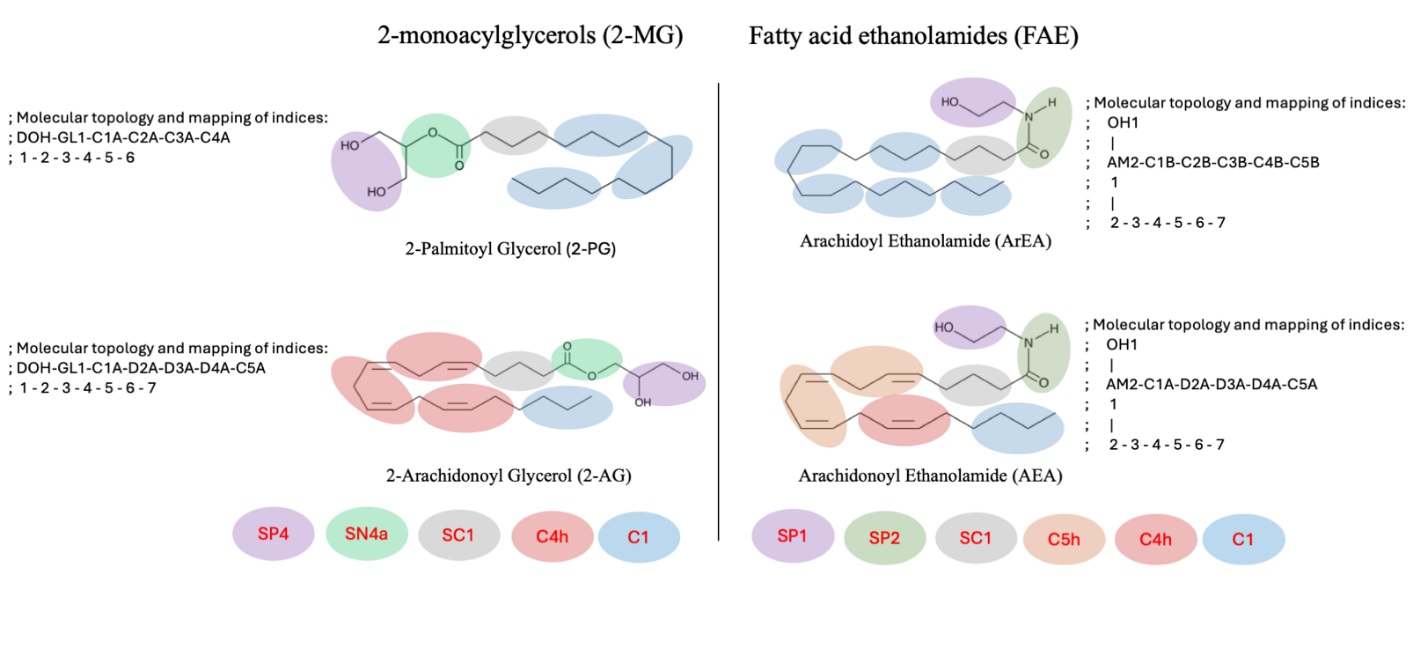
